# A two-state ribosome and protein model can robustly capture the chemical reaction dynamics of gene expression

**DOI:** 10.1101/2020.11.25.399287

**Authors:** Ayush Pandey, Richard M. Murray

**Affiliations:** Control and Dynamical Systems, California Institute of Technology, Pasadena, CA, 91125; Biology and Biological Engineering, California Institute of Technology, Pasadena, CA, 91125

## Abstract

We derive phenomenological models of gene expression from a mechanistic description of chemical reactions using an automated model reduction method. Using this method, we get analytical descriptions and computational performance guarantees to compare the reduced dynamics with the full models. We develop a new two-state model with the dynamics of the available free ribosomes in the system and the protein concentration. We show that this new two-state model captures the detailed mass-action kinetics of the chemical reaction network under various biologically plausible conditions on model parameters. On comparing the performance of this model with the commonly used mRNA transcript-protein dynamical model for gene expression, we analytically show that the free ribosome and protein model has superior error and robustness performance.

## 1 Background

For model-based design of biological circuits, we need to develop mathematical models that map the system design specifications to the mechanistic details. Commonly used phenomenological models are based on empirical information and their model parameters describe lumped properties of the system that are effective in explaining the observed experimental data [1–5] but have not been readily used for forward engineering of biological circuits. Towards that end, to explore different design possibilities one needs to carefully justify the validity of the underlying assumptions for each model [6]. This is demonstrated in a simple analysis by Del Vecchio and Murray [7] for an enzymatic reaction system. It is shown that the commonly used Michaelis-Menten [8] kinetics for the enzymatic reaction system fails to capture the true dynamics under different parameter regimes. In this paper, we discuss a particular example of the expression of a single gene, and analytically derive phenomenological models with exactly known mappings to the mechanistic details. We explore the modeling assumptions of time-scale separation [9–11], conservation laws [7, 12, 13] and prove the robustness of various models under different parametric conditions.

For gene expression, a two-state model is commonly used in the literature [1, 2, 14–16] that models the dynamics of the mRNA transcript (*T*) and the protein concentration (*X*) as a function of the DNA copy number (*G*) and regulatory effects:

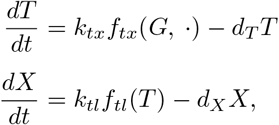

where *k*_*tx*_ is defined as the transcription rate and *k*_*tl*_ is the translation rate. Similarly, *d*_*T*_ and *d*_*X*_ are the degradation and dilution parameters for the transcript and the protein respectively. The function *f*_*tx*_(·) is usually a Hill function dependent on the mechanism of transcription-factor activation or repression. For constitutive expression, this is assumed to be a constant function of the DNA copy number, *f*_*tx*_(*G*) = *kG*. Similarly, *f*_*tl*_(·) could be a constant or a Hill function dependent on the transcriptional regulation mechanism [7]. Clearly, the parameters in this model and any parameters in the Hill functions all have empirical meanings but an analytical relationship with the mechanistic reaction rates is usually obscured [17]. Moreover, a closer analysis would show that such phenomenological models are only valid under certain assumptions and parameter regimes. In this paper, we use an automated model reduction approach [18] to develop reduced-order models for the gene expression example. We give an analytical characterization of the lumped parameters to the mechanistic details along with a clear description of various underlying assumptions and performance guarantees. Our main result shows that for gene expression a new two-state model of the free ribosomes and protein dynamics can capture the detailed chemical reaction network dynamics under a wide range of parameter values.

## 2 Mathematical models of gene expression

### 2.1 The full CRN model

The full chemical reaction network (CRN) model for the expression of protein *X* from a single gene *G* is described in Table 1. In this CRN, the gene *G* is transcribed by RNA polymerase (*P*) to an mRNA transcript *T* via a complex (*C*_1_) formation reaction. Then, the transcript *T* binds to the ribosome *R* to form the second complex *C*_2_, that then translates to express the protein *X*. Under the assumption of mass-action kinetics for all reactions, the ordinary differential equation (ODE) model can be derived as shown on the right in Table 1. We refer to this as the full CRN model for the rest of this paper.

**Table 1:** Full CRN and the corresponding ODE model assuming mass-action kinetics obtained using a CRN compiler software [19].

| CRN | ODE model |
| --- | --- |
| $G + P \xrightleftharpoons[k_{up}]{k_{bp}} C_1$ | $\frac{dP}{dt} = (k_{up} + k_{tx}) C_1 - k_{bp} GP$ |
| $C_1 \xrightarrow{k_{tx}} G + P + T$ | $\frac{dC_1}{dt} = k_{bp} GP - (k_{up} + k_{tx}) C_1$ |
| $T + R \xrightleftharpoons[k_{ur}]{k_{br}} C_2$ | $\frac{dT}{dt} = k_{tx} C_1 + (k_{ur} + k_{tl}) C_2$<br>$- k_{br} TR - d_T T$ |
| $C_2 \xrightarrow{k_{tl}} T + R + X$ | $\frac{dR}{dt} = (k_{ur} + k_{tl}) C_2 - k_{br} TR$ |
| $T \xrightarrow{d_T} \emptyset$ | $\frac{dC_2}{dt} = k_{br} TR - (k_{ur} + k_{tl}) C_2$ |
| $X \xrightarrow{d_X} \emptyset$ | $\frac{dX}{dt} = k_{tl} C_2 - d_X X$ |

### 2.2 Reduced-order modeling

Model reduction [18, 20–22] is a widely used tool in engineering design and analysis. Abstracting away the details of a system model to focus on modeling the properties of interest and its interactions is an important insight that is commonly used in control systems’ design. Reduced models are also useful to specify the desired objectives or the performance specifications of a system. To meet these objectives, the designer needs to map these reduced models to the level of system design and also mathematically characterize this mapping in order to analyze the system performance [17, 23]. Towards that end, it is important to explore the modeling assumptions of time-scale separation [9–11] and conservation laws [7, 12, 13] that are most commonly used in deriving reduced-order models for biological systems. We study this using an example of gene expression. We analyze the robustness of various gene expression models under different parametric conditions. Using this analysis, we propose a forward design approach to guide the design of synthetic biological circuits.

To obtain all reduced model expressions we developed an automated model reduction software [24], `AutoReduce`. This software can solve for conservation laws to derive reduced order models symbolically. More importantly, we can automatically obtain all possible reduced models under time-scale separation assumptions. Time-scale separation is one of the most commonly used assumptions in biological systems where a set of species reaches steady-state quickly and so the dynamics of the slow set of species can be approximated by assuming the fast species at quasi-steady state (QSS). However, it is important to perform this dynamics reduction carefully as the reduced models may or may not capture the full-order model dynamics under different parameter regimes.

For the gene expression chemical reaction network, the first step is to solve for the conserved quantities in the model. We assume that the total RNA polymerase in the system and the total ribsomes remain conserved. From Table 1 observe that

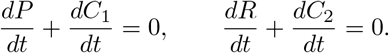

Hence, for constants *P*_tot_ and *R*_tot_, we have that *P*_tot_ = *P* + *C*_1_ and *R*_tot_ = *R* + *C*_2_. These conservation laws can be used to eliminate *C*_1_ and *C*_2_. Using `AutoReduce`, we obtained the following reduced-order model under conservation laws,

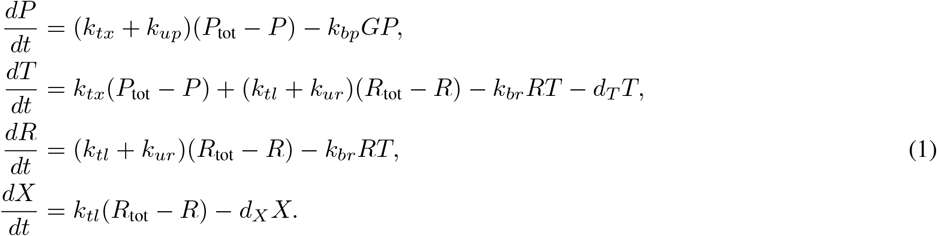

Next, we obtain various reduced models under different time-scale separation assumptions. We can analytically derive these reduced models using `AutoReduce` and explore their validity and performance guarantees. We denote all reduced-order model variables with a hat to differentiate the corresponding variable in the full model, for example, in a reduced model the protein species will be represented as 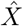 and the corresponding species in the full model is denoted by *X*. All variables denote the concentrations for each species and parameters take appropriate units. Furthermore, we define the following lumped parameter notations which appear as Hill function activation parameters in the reduced model expressions that we derive next.

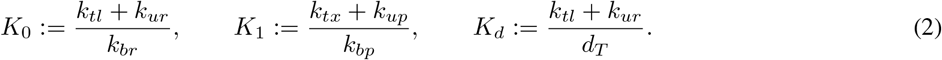

## 3 Results

### 3.1 Simple gene expression

We start by presenting various reduced-order models for the simple gene expression chemical reaction network and the corresponding ODE model is given in Table 1.

#### The mRNA transcript 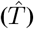 and protein 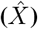 model

Under the assumption that the free ribosomes and the RNA polymerase dynamics are at QSS, we obtain the following model with only the mRNA transcript and the protein dynamics as a function of the DNA copy number *G*:

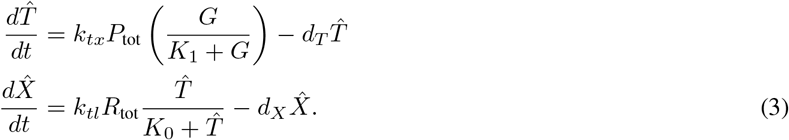

#### The free ribosome 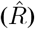 and protein 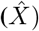 model

Under the assumption that the mRNA transcript and the RNA polymerase dynamics are at QSS, we obtain the following model with only the free ribosome and the protein dynamics:

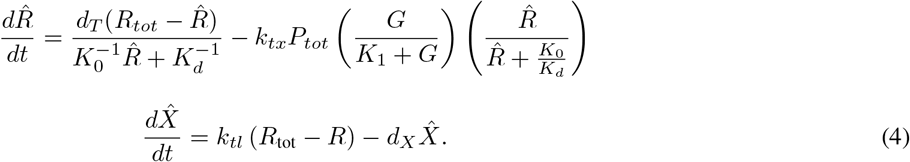

Similar to the ribosome-protein and the mRNA transcript-protein models, it is possible to derive the polymerase-protein ([*P, X*]) and the only protein model ([*X*]). The detailed equations for these two models are not presented here for brevity but their performance is shown in Figures 1 and 2.

**Figure 1:**
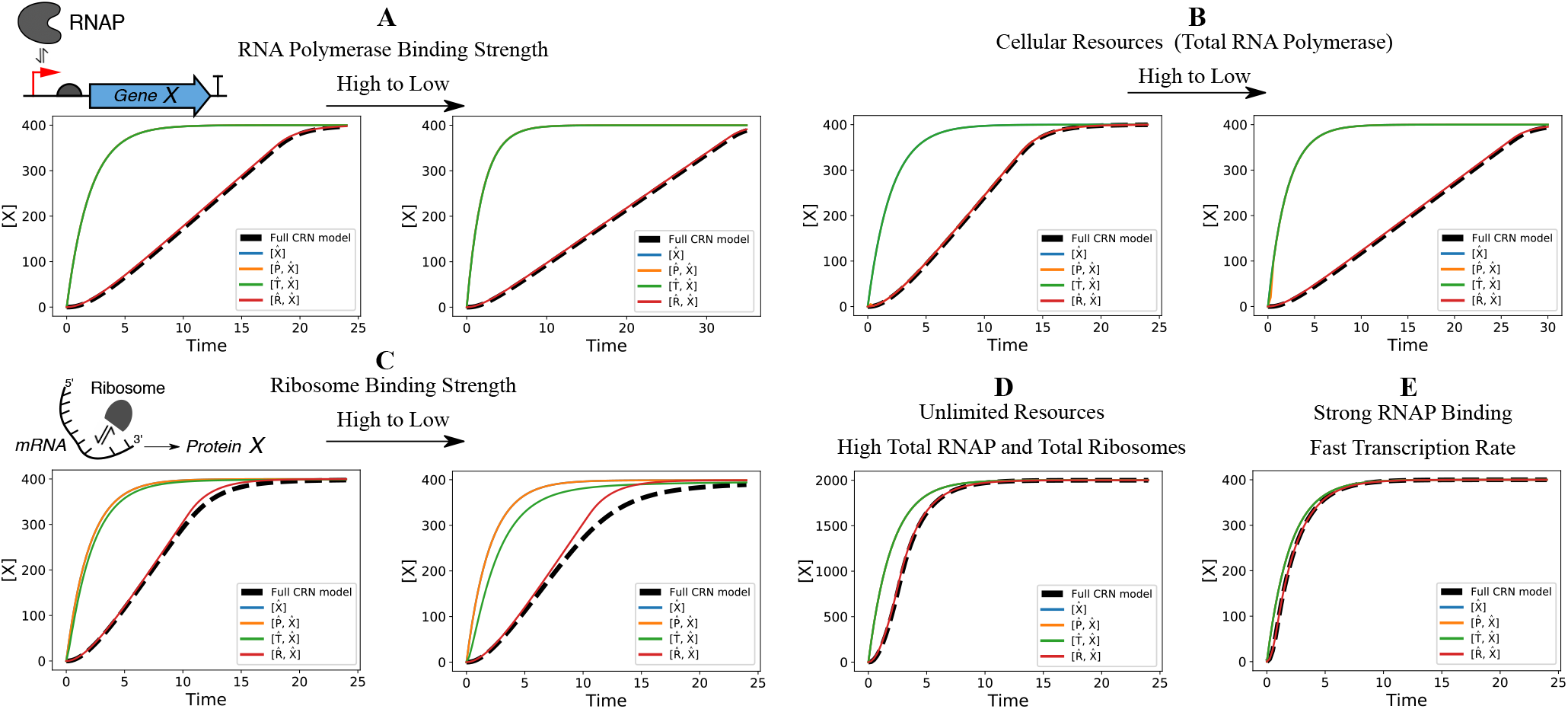
Performance of the gene expression models under different biological conditions. **(A)** We observe that for weaker binding of RNA polymerase to the promoter region of the DNA, the time-response of the full CRN model is slower as it takes a longer time to reach steady-state. The mathematical model with only the mRNA transcript and protein dynamics is unable to capture this effect since this binding reaction is assumed to be at quasi-steady state in this model. On the other hand, the *R*_Δ_ model, which describes the dynamics of the free ribosome and the protein is able to capture the effect. **(B)** With decreasing total RNA polymerase, the time-response of the full model is slower and only the ribosome-protein model is able to account for this effect. **(C)** With decreasing ribosome binding strength, none of the reduced models perfectly capture the full CRN dynamics, but still, the ribosome-protein model is the closest in error performance to the full model. **(D)** With a very high amount of total RNA polymerase and the total ribosome count in the system, all models reach steady-state faster. **(E)** Under strong RNA polymerase binding to the DNA and high transcription rate, we see that all models exhibit good error performance. These can be understood as the ideal conditions under which using a one-state protein dynamics model is also justified. Python code used for this analysis is available publicly on Github [25] and can also be run online using this link.

**Figure 2:**
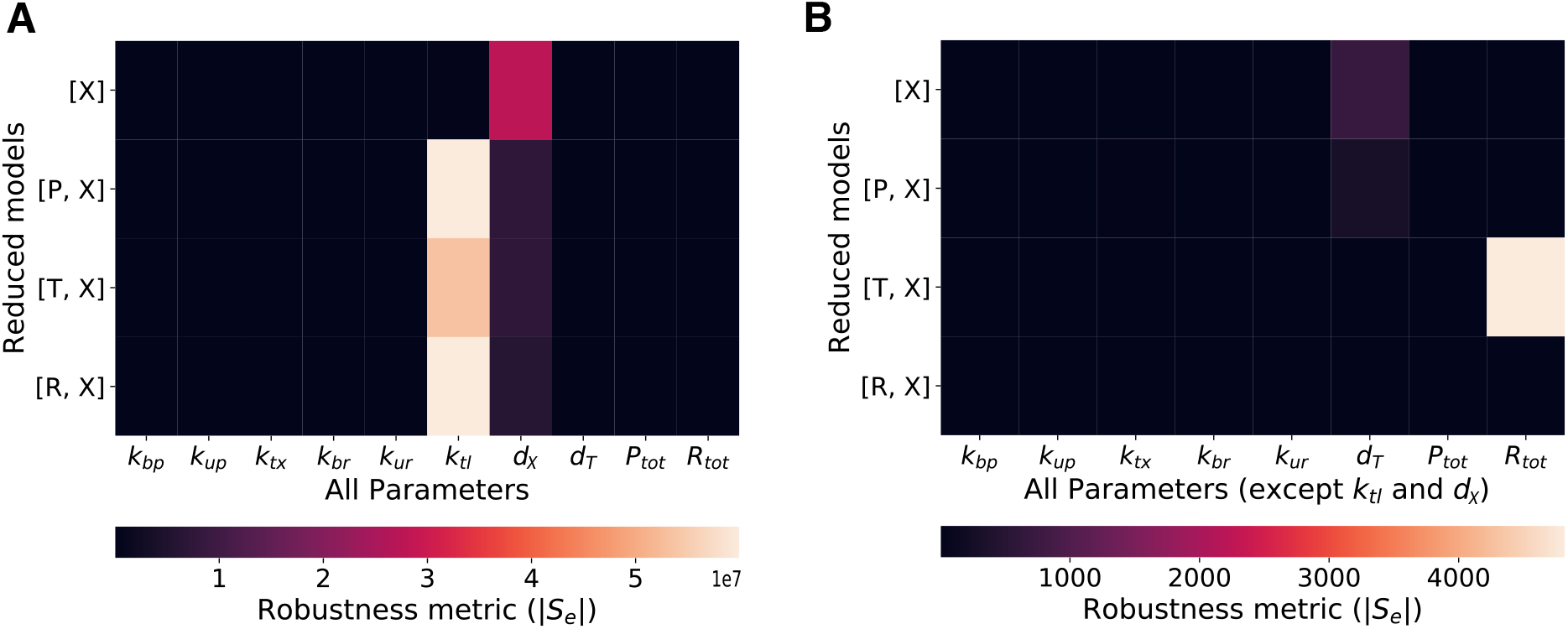
The figure shows the fragility of different models with respect to all model parameters as computed by the sensitivity of the error in protein *X* concentration between the full model and the reduced models. **(A)** All reduced models are fragile to the translation parameter *k*_*tl*_, and the protein degradation parameter *d*_*X*_, as shown analytically in Statement 1. **(B)** Other than *k*_*tl*_ and *d*_*X*_, the performance of the *R*_Δ_ model (model with free ribosome and protein dynamics) is not fragile to any other model parameters, however, other models, in particular the commonly used mRNA transcript and protein dynamical model is not robust to the total ribosome count *R*_tot_. Python code used for this analysis is available publicly on Github [25].

#### The available free ribosome 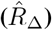 and protein 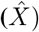 model

We define the available free ribosomes in the ge[ne expre]ssion system as *R*_Δ_ = *R*_tot_ – *R*. Substituting 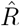 for 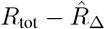, we can derive a new reduced-order model from the 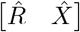 model – the *R*_Δ_ model:

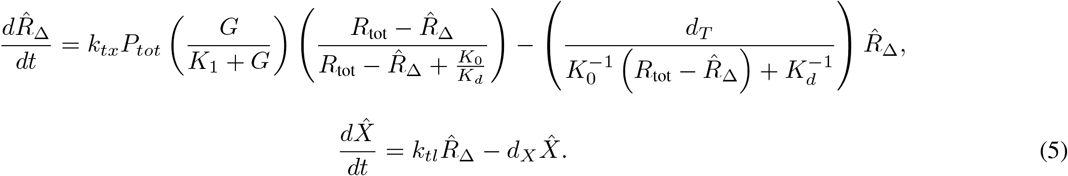

This model simplifies the robustness analysis that follows. Note that the *R*_Δ_ model closely resembles the commonly used gene expression model and all of its terms are exactly similar to the mRNA transcript and protein model but scaled by a ribosome count factor.

##### Statement 1.

*The error between the R*_Δ_ *model and the full* ***CRN*** *model is robust to perturbations in the transcription rate* (*k*_*tx*_), *binding/unbinding of polymerase* (*k*_*bp*_, *k*_*up*_) *and ribosome* (*k*_*br*_, *k*_*ur*_), *and also total resources (both RNA polymerase, P*_*tot*_ *and ribosome, R*_*tot*_). *Only the perturbations in the translation rate* (*k*_*tl*_) *and the protein degradation* (*d*_*X*_) *parameters exacerbate the error performance of the R*_Δ_ *model*.

*Justification*. To demonstrate robustness of the *R*_Δ_ model, we can look at the sensitivity of the error between the *R*_Δ_ model and the full CRN model to various parameter perturbations. Define the error as 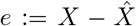, where *X* and 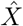 represent the protein concentration in the full CRN model and the *R*_Δ_ model respectively. In the case where the error *e* between the two models is within acceptable bounds, we additionally desire that this error is not sensitive to any of the parameters. The sensitivity of the error to perturbation in a parameter *θ* is given by *S*_*e*_ = *∂e/∂θ*. Hence, this “fragility metric” *S*_*e*_ must be minimized to achieve higher robustness. Analyzing the rate of change of *S*_*e*_ with time for different parameters can justify the statement above.

To analytically derive *Ṡ*_*e*_, we utilize the results for the sensitivity system equation [18]. In summary, to derive *Ṡ*_*e*_ for any parameter *θ*, we first solve for the Jacobian matrices 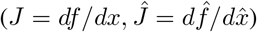 for both the full model, 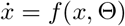 and the reduced model 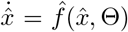, where *x*, 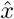 represent the model states and Θ is the set of model parameters. Note that the model output is the protein concentration, given in matrix terms as *y* = *Cx* and 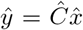 where *C* and 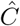 are row matrices such that we have *y* = *X* and 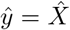, the protein concentrations. We also solve for the sensitivity to parameter *Z* = *df/dθ* for both models. Finally, *Ṡ*_*e*_ is given by [18],

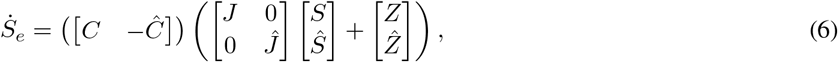

where *S* = *dx/dθ* and 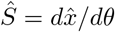 are the sensitivity coefficients of the full model and the reduced model respectively.

For the parameters *k*_*tx*_, *k*_*bp*_, *k*_*up*_, *k*_*br*_, *k*_*ur*_, *P*_tot_, and *R*_tot_, we have the dynamics of fragility metric *S*_*e*_ given by

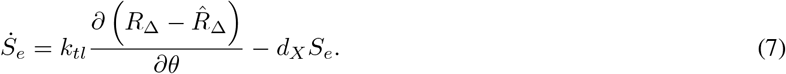

For the protein degradation rate *d*_*X*_, we have

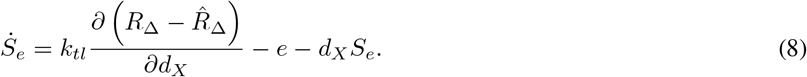

For the translation rate *k*_*tl*_, we have

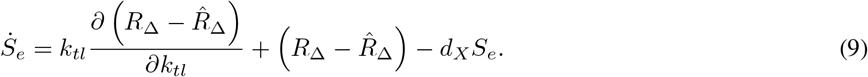

From equations (7) – (9), we have that for all positive parameter values and stable models, the dynamics of *S*_*e*_ are convergent to a fixed point. More importantly, the fragility metric, *S*_*e*_ directly depends on the translation rate *k*_*tl*_ and the protein degradation rate *d*_*X*_, proving the assertion of the statement above. Also, for *k*_*tl*_ and *d*_*X*_, we see that an extra term appears in the error sensitivity dynamics implying higher fragility of the *R*_Δ_ model under perturbations to the translation rate and the protein degradation rate.

Figure 1 shows the error performance of the *R*_Δ_ model under various biologically plausible parameter conditions and assumptions. The Euclidean norm of the fragility metric, *S*_*e*_, is plotted in Figure 2 to compare the robustness of the *R*_Δ_ model alongside other models. Similar to the result above, we give the following statement for the transcript and the protein model.

##### Statement 2.

*The mathematical model with mRNA transcript and protein dynamics, given in equation (3), captures the full CRN model dynamics only under the assumption of unlimited resources and fast binding reactions. As a result, the error performance of the transcript-protein model is directly dependent on the mRNA-ribosome binding/unbinding parameters and the translation parameter. Hence, unlike the R*_Δ_ *model, this model is not robust to perturbations in k*_*br*_, *k*_*ur*_, *and R*_*tot*_ *in addition to k*_*tl*_ *and d*_*X*_.

*Justification.* Similar to the proof of Statement 1, we can analyze the dynamics of the sensitivity of the error to parameter perturbations for the mRNA transcript and protein dynamical model.

For the translation rate, that is *θ* = *k*_*tl*_, we have,

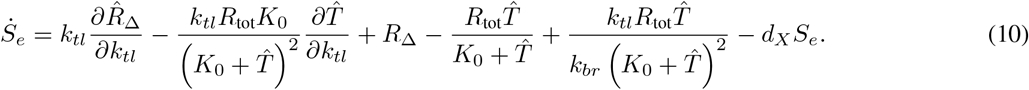

For the protein degradation parameter *θ* = *d*_*X*_, we have,

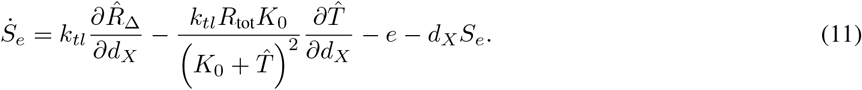

For the total ribosome count, *θ* = *R*_tot_, we have,

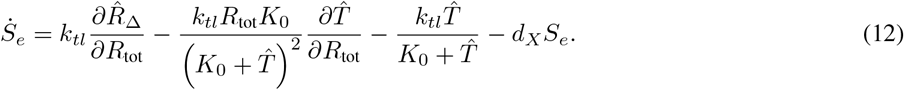

For the ribsome-transcript binding parameter, *θ* = *k*_*br*_, we have,

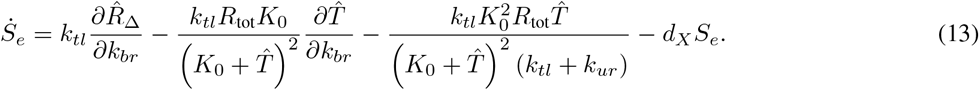

For the ribsome-transcript unbinding parameter, *θ* = *k*_*ur*_, we have,

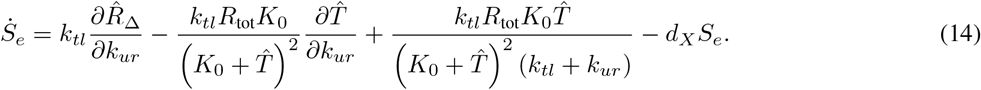

For all other parameters, we have,

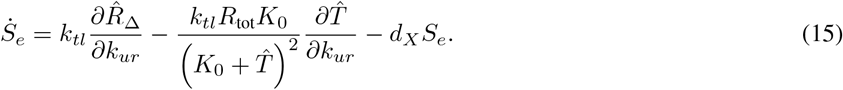

From equations (10) – (15), we can conclude that the fragility metric, *S*_*e*_, directly depends on the total ribosome count, *R*_tot_, the binding/unbinding parameters of the transcript with ribosome, *k*_*br*_, *k*_*ur*_, the translation rate, *k*_*tl*_, and the protein degradation rate, *d_X_*. This is in contrast with the results for the *R*_Δ_ model where the fragility only depends on the translation rate and the protein degradation rate. Hence, the *R*_Δ_ model is robust to all other parameters whereas the transcript-protein model is fragile to all of these parameters, proving the statement assertion. Figure 2 numerically shows this result.

### 3.2 Gene expression with endonuclease mediated mRNA degradation

To investigate the scalability of this approach, we expand on the gene expression example by including enzymatic degradation of the mRNA transcript mediated by endonucleases. Here we have this enzymatic degradation in addition to the basal degradation rate *d_T_*. The CRN and the corresponding mass-action ODE model is given in Table 2.

**Table 2:** Full CRN model for gene expression with endonuclease mediated mRNA degradation

| CRN | ODE model |
| --- | --- |
| $G + P \xrightleftharpoons[k_{up}]{k_{bp}} C_1$ | $\frac{dP}{dt} = (k_{up} + k_{tx}) C_1 - k_{bp} GP$ |
| $C_1 \xrightarrow{k_{tx}} G + P + T$ | $\frac{dC_1}{dt} = k_{bp} GP - (k_{up} + k_{tx}) C_1$ |
| $T + R \xrightleftharpoons[k_{ur}]{k_{br}} C_2$ | $\frac{dT}{dt} = k_{tx} C_1 + (k_{ur} + k_{tl}) C_2 - k_{br} TR + k_{ue} C_3 - k_{be} TE - d_T T$ |
| $C_2 \xrightarrow{k_{tl}} T + R + X$ | $\frac{dR}{dt} = (k_{ur} + k_{tl}) C_2 - k_{br} TR$ |
| $T + E \xrightleftharpoons[k_{ue}]{k_{be}} C_3$ | $\frac{dC_2}{dt} = k_{br} TR - (k_{ur} + k_{tl}) C_2$ |
| $C_3 \xrightarrow{d_E} E$ | $\frac{dE}{dt} = (k_{ui} + d_E) C_3 - k_{be} TE$ |
| | $\frac{dC_3}{dt} = k_{be} TE - (k_{ue} + d_E) C_3$ |
| $X \xrightarrow{d} \emptyset$ | $\frac{dX}{dt} = k_{tl} C_2 - d_X X$ |

Observe that in this model we have the following conservation law relationships,

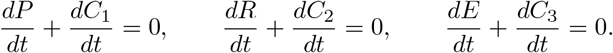

Hence, for constants *P*_tot_, *R*_tot_, and *E*_tot_, we can write

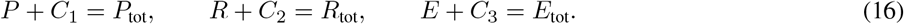

Using these algebraic relationships, we can eliminate the complexes *C*_1_, *C*_2_, and *C*_3_ to obtain a reduced ODE model. Next, we explore various time-scale separation assumptions that may be used to get further reduced models and discuss their performance and robustness with respect to the model parameters.

#### The mRNA transcript and protein dynamical model

Assuming that the dynamics of all species in the model other than the mRNA transcript *T* and the protein *X* are at quasi-steady state, we get the following model. Observe that a new degradation Hill function term appears in the dynamics of the mRNA transcript that is dependent on the endonuclease binding parameters.

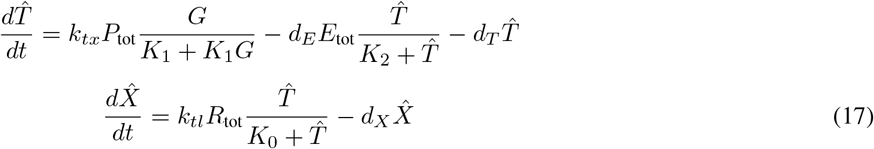

where *K*_2_ is a new lumped parameter that is the Hill activation parameter for the endonuclease binding,

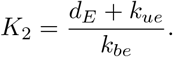

#### Modeling the dynamics of available free ribosomes

As before, we define the available free ribosomes in the system as *R*_Δ_ = *R*_tot_ – *R*. Using this definition and assuming that the dynamics of all species in the model other than the free ribosomes, the mRNA transcript, and the protein are at quasi-steady state, we get the following model:

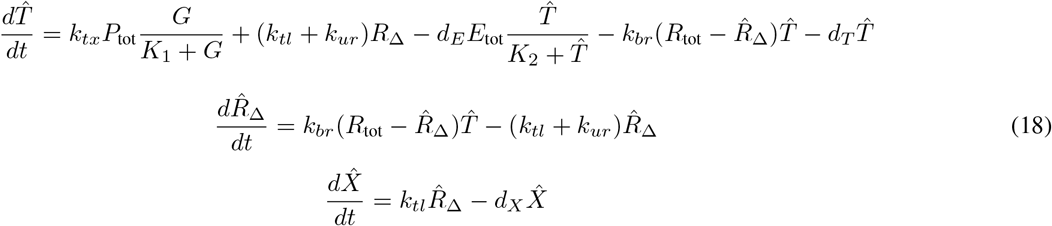

Note that the model with only *R*_Δ_ and the protein *X* dynamics does not work in this case where the enzymatic degradation reactions for the mRNA transcript are significant for the overall dynamics. So, either *T* or *E* is necessary in the ribosome-protein model to get satisfactory performance. As a result, we can also obtain a reduced model with 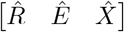 as the states. The performance of all of the possible reduced models is shown in Figure 3.

**Figure 3:**
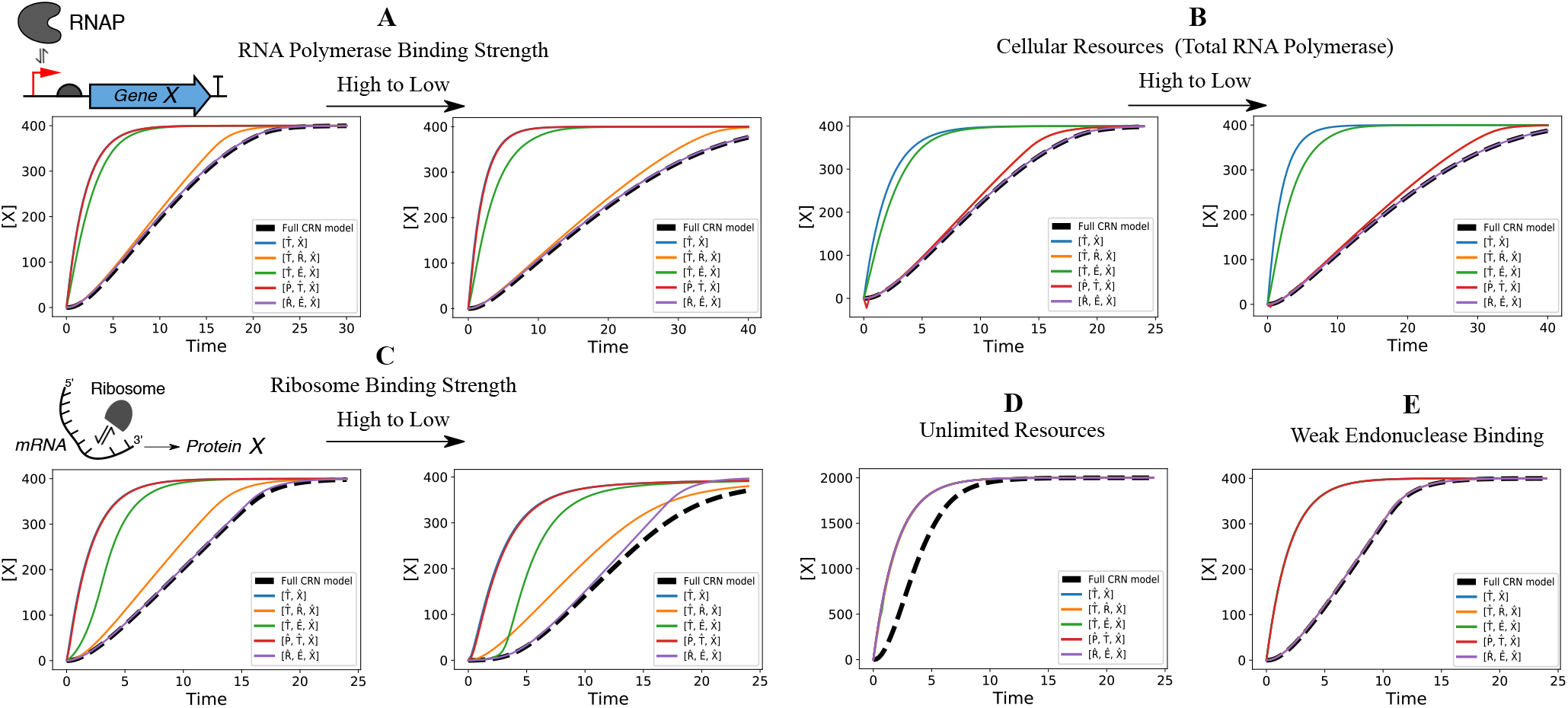
Performance of the gene expression with endonuclease mediated mRNA degradation models under different biological conditions. **(A)** Similar to Figure 1, we observe that for weaker binding of RNA polymerase to the promoter region of the DNA implies a slower time-response which is only captured perfectly by the free ribosome model that also models the endonuclease dynamics. The mathematical model with only the mRNA transcript and protein dynamics is unable to capture this effect since this binding reaction is assumed to be at quasi-steady state in this model, but it is the only two-state model for this system with a satisfactory steady-state performance. **(B)** We observe a similar effect with decreasing total RNA polymerase in the system. **(C)** With decreasing ribosome binding strength, none of the reduced models perfectly capture the full CRN dynamics, but still, the ribosome-protein model with endonuclease dynamics is the closest in error performance to the full model. **(D)** With a very high amount of cellular resources, all models reach steady-state faster. **(E)** Under weak endonuclease binding, we observe that only the model that explicitly models the endonuclease dynamics gives good performance. Python code used for this analysis is available publicly on Github [25].

Although a similar robustness analysis as shown in Statements 1 and 2 can be done for this system dynamics it would be easier to numerically compute the fragility metric *S*_*e*_ for all reduced models. The heatmap of the Euclidean norm of this metric is shown in Figure 4. To exactly compute this metric, all sensitivity coefficients for all reduced models and the full model would have to be computed. This would be computationally inefficient for a higher number of states in the models. So, we use an upper bound to *S*_*e*_ derived in [18] to compute the fragility metric. The `AutoReduce` software can be used to compute the model equations and these metrics and so a future line of work would be to extend this approach to a realistic circuit design with multiple gene expression components.

**Figure 4:**
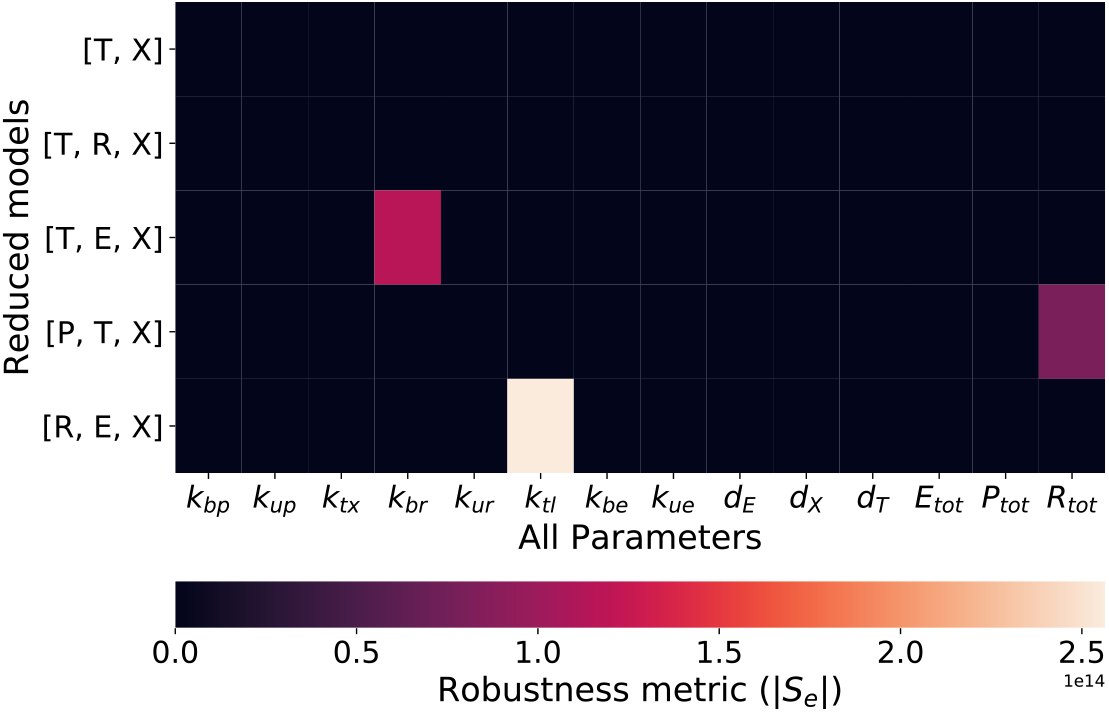
The figure shows the fragility of different models with respect to all model parameters as computed by the sensitivity of the error in protein *X* concentration between the full model and the reduced models.

## 4 Discussion

The exploration of design space by changing experimentally tunable parameters in the models as shown in Figure 1 is an important step towards using mathematical models for biological circuit design. There are two main advantages to this approach:

### 4.1 Circuit design

For any complex biological circuit, a single gene expression would usually form a small part of the design. Hence, inter-preting the key states and parameters involved in tuning its dynamics is important for the modular design of the complete circuit. This task becomes increasingly hard if full CRN descriptions were used for each module instead. So, reduced-order models for each module with a known mapping to the full dynamics could assist in a model-based design approach. Towards that end, a future line of work could be to develop a pipeline that is compatible with the CRN compiler software called BioCRNpyler [19] so that we can obtain reduced models for CRNs of different modules in a circuit. A potential application of this approach to circuit design is discussed in Figure 5.

**Figure 5:**
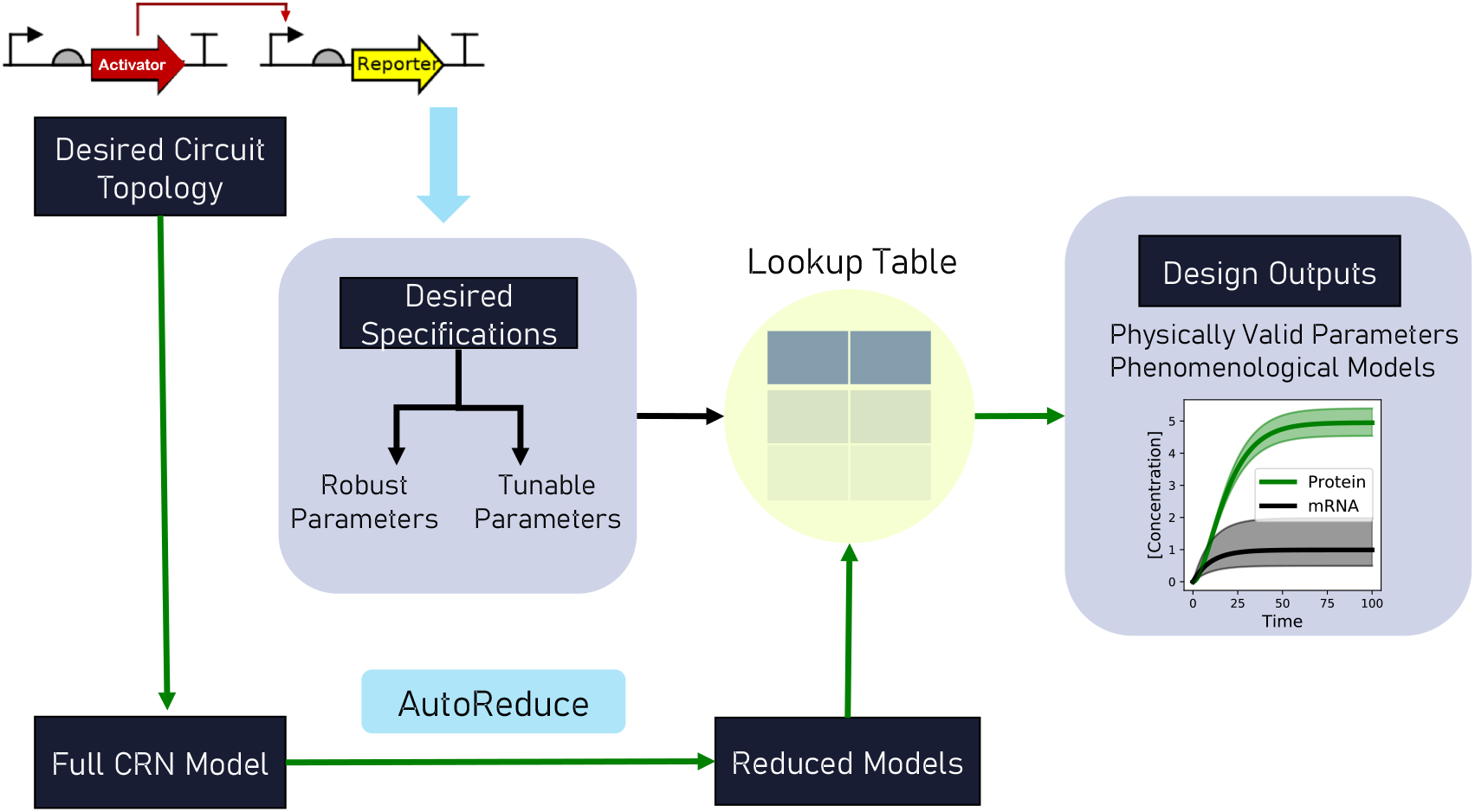
The figure shows a possible forward design pipeline for synthetic biological circuits. Given a desired circuit topology, the user can specify a list of specifications for the circuit design dependent on the objectives. These specifications could be in the form of a numerical range for the tunable parameters in the experiment – for example, the ribosome binding strength. The user could also specify the parameters that the experiment design should be robust to – for example, total resources. Given the specifications, the challenge is to give physically valid parameter ranges for the tunable parameters and a dynamical model that can be used for prediction and analysis. The path with the green arrows in the figure is the proposed automated pipeline for this forward design approach. Using an automated software like BioCRNpyler [19], a full CRN model can be created given the circuit topology. Then, using `AutoReduce`, we can obtain reduced-order models and create a lookup table using analysis similar to the results in Figures 1 and 3. Using this lookup table, we could give design outputs in terms of valid parameter ranges for the tuned parameters and guarantees under bounded parametric uncertainties for other model parameters as shown in the rightmost block in the figure. A reduced model describing the effective outputs is also available as an output that is helpful for system analysis and parameter inference.

### 4.2 System analysis and parameter inference

Empirical reduced-order models using Hill functions are already commonplace in the literature for parameter identification of biological circuits. However, as discussed through the example of gene expression, models are only valid as long as the underlying assumptions are biologically valid. Hence, it is important to use the correct reduced-order model for parameter inference and system analysis. The automated model reduction method discussed in this paper is a step towards that goal as it provides a mapping of the reduced models with the full model as well as performance guarantees to verify if a reduced model is valid under certain assumptions or not.

In our model reduction approach, all lumped parameter relationships are known. Due to parameter lumping, the reduced models have fewer number of parameters, improving the parameter identifiability of the system given measurement data [26]. Moreover, the non-identifiable manifolds are analytically known as well. Hence, reduced models with known mapping to the full models are helpful in the parameter inference problem, especially for synthetic biological systems where only a few measurements are available.

An interesting future direction would be to extend the results in this paper for expression of multiple genes together to explore retroactivity [27] and its effects on various phenomenological models used for design of such systems. This study would be a step in building towards a modular design framework [28] that considers the design of multiple modules together with their context-dependence. Representing such huge models in phenomenological terms is a challenging task with the current model reduction approach. Hence, derivation of reduced models by introducing defined coordinate transformations for states and parameters might be worth investigating as well.

## Acknowledgement

The authors thank Ankita Roychoudhury for improving and providing feedback on the `AutoReduce` Python package that is used to derive the different models shown in this document. This research is sponsored in part by the National Science Foundation under grant number: CBET-1903477 and the Defense Advanced Research Projects Agency (Agreement HR0011-17-2-0008). The content of the information does not necessarily reflect the position or the policy of the Government, and no official endorsement should be inferred.

## References

[1] Michael B Elowitz and Stanislas Leibler. “A synthetic oscillatory network of transcriptional regulators”. In: Nature 403.6767 (2000), pp. 335–338.

[2] Timothy S Gardner, Charles R Cantor, and James J Collins. “Construction of a genetic toggle switch in Escherichia coli”. In: Nature 403.6767 (2000), pp. 339–342.

[3] Chelsea Y Hu, Jeffrey D Varner, and Julius B Lucks. “Generating effective models and parameters for RNA genetic circuits”. In: ACS synthetic biology 4.8 (2015), pp. 914–926.

[4] Reed D. McCardell, Ayush Pandey, and Richard M. Murray. “Control of density and composition in an engineered two-member bacterial community”. In: bioRxiv (2019). DOI: 10.1101/632174.

[5] Anandh Swaminathan, Victoria Hsiao, and Richard M Murray. “Quantitative modeling of integrase dynamics using a novel python toolbox for parameter inference in synthetic biology”. In: bioRxiv (2017), p. 121152.

[6] Andras Gyorgy and Domitilla Del Vecchio. “Limitations and trade-offs in gene expression due to competition for shared cellular resources”. In: 53rd IEEE Conference on Decision and Control. IEEE. 2014, pp. 5431–5436.

[7] Domitilla Del Vecchio and Richard M Murray. Biomolecular Feedback Systems. Princeton University Press Princeton, NJ, 2015.

[8] Kenneth A Johnson and Roger S Goody. “The original Michaelis constant: translation of the 1913 Michaelis–Menten paper”. In: Biochemistry 50.39 (2011), pp. 8264–8269.

[9] Andrei Nikolaevich Tikhonov. “Systems of differential equations containing small parameters in the derivatives”. In: Matematicheskii Sbornik 73.3 (1952), pp. 575–586.

[10] Alexandra Goeke, Sebastian Walcher, and Eva Zerz. “Determining “small parameters” for quasi-steady state”. In: Journal of Differential Equations 259.3 (2015), pp. 1149–1180.

[11] Thomas J Snowden, Piet H van der Graaf, and Marcus J Tindall. “Methods of model reduction for large-scale biological systems: a survey of current methods and trends”. In: Bulletin of mathematical biology 79.7 (2017), pp. 1449–1486.

[12] Thomas J Snowden, Piet H Van Der Graaf, and Marcus J Tindall. “A combined model reduction algorithm for controlled biochemical systems”. In: BMC Systems Biology 11.1 (2017), p. 17.

[13] Manvel Gasparyan, Arnout Van Messem, and Shodhan Rao. “An Automated Model Reduction Method for Biochemical Reaction Networks”. In: Symmetry 12.8 (2020), p. 1321.

[14] Olguta Buse, Rodrigo Pérez, and Alexey Kuznetsov. “Dynamical properties of the repressilator model”. In: Physical Review E 81.6 (2010), p. 066206.

[15] Chelsea Y Hu and Richard M Murray. “Design of a genetic layered feedback controller in synthetic biological circuitry”. In: bioRxiv (2019), p. 647057.

[16] Adam J Meyer et al. “Escherichia coli “Marionette” strains with 12 highly optimized small-molecule sensors”. In: Nature chemical biology 15.2 (2019), pp. 196–204.

[17] Lorenzo Pasotti et al. “Mechanistic models of inducible synthetic circuits for joint description of DNA copy number, regulatory protein level, and cell load”. In: Processes 7.3 (2019), p. 119.

[18] Ayush Pandey and Richard M Murray. “An automated model reduction tool to guide the design and analysis of synthetic biological circuits”. In: bioRxiv (2019), p. 640276.

[19] William Poole et al. “BioCRNpyler: Compiling Chemical Reaction Networks from Biomolecular Parts in Diverse Contexts”. In: bioRxiv (2020).

[20] Petar V Kokotovic, Robert E O’Malley Jr, and Peddapullaiah Sannuti. “Singular perturbations and order reduction in control theory - an overview”. In: Automatica 12.2 (1976), pp. 123–132.

[21] Bruce Moore. “Principal component analysis in linear systems: Controllability, observability, and model reduction”. In: IEEE Transactions on Automatic Control 26.1 (1981), pp. 17–32.

[22] Antonis Papachristodoulou et al. “Structured model reduction for dynamical networked systems”. In: 49th IEEE Conference on Decision and Control (CDC). IEEE. 2010, pp. 2670–2675.

[23] Mark K Transtrum and Peng Qiu. “Bridging mechanistic and phenomenological models of complex biological systems”. In: PLoS computational biology 12.5 (2016), e1004915.

[24] Ayush Pandey and Richard M. Murray. “Model Reduction Tools For Phenomenological Modeling of Input-Controlled Biological Circuits”. In: bioRxiv (2020). DOI: 10.1101/2020.02.15.950840.

[25] Ayush Pandey. Auto-Reduce: Python based automated model reduction package. [Online].

[26] Florin Paul Davidescu and Sten Bay Jørgensen. “Structural parameter identifiability analysis for dynamic reaction networks”. In: Chemical Engineering Science 63.19 (2008), pp. 4754–4762.

[27] Shridhar Jayanthi, Kayzad Soli Nilgiriwala, and Domitilla Del Vecchio. “Retroactivity controls the temporal dynamics of gene transcription”. In: ACS synthetic biology 2.8 (2013), pp. 431–441.

[28] Andras Gyorgy and Domitilla Del Vecchio. “Modular composition of gene transcription networks”. In: PLoS Comput Biol 10.3 (2014), e1003486.

